# An integrative data-centric approach to derivation and characterization of an adverse outcome pathway network for cadmium-induced toxicity

**DOI:** 10.1101/2023.12.12.571252

**Authors:** Ajaya Kumar Sahoo, Nikhil Chivukula, Kundhanathan Ramesh, Jasmine Singha, Shambanagouda Rudragouda Marigoudar, Krishna Venkatarama Sharma, Areejit Samal

**Author notes:** Corresponding author (A. Samal); Address for correspondence: Areejit Samal, Computational Biology Group, The Institute of Mathematical Sciences (IMSc), CIT Campus, Taramani, Chennai 600113 India. A.K.S. and N.C. contributed equally to this work and should be considered as Joint-First authors.

## Abstract

Cadmium is a prominent toxic heavy metal that contaminates both terrestrial and aquatic environments. Owing to its high biological half-life and low excretion rates, cadmium causes a variety of adverse biological outcomes. Adverse outcome pathway (AOP) networks were envisioned to systematically capture toxicological information to enable risk assessment and chemical regulation. Here, we leveraged AOP-Wiki and integrated heterogeneous data from four other exposome-relevant resources to build the first AOP network relevant for inorganic cadmium-induced toxicity. From AOP-Wiki, we filtered 309 high confidence AOPs, identified 312 key events (KEs) associated with inorganic cadmium, and thereafter, curated 30 cadmium relevant AOPs (cadmium-AOPs), using a data-centric approach. By constructing the undirected AOP network, we identified a large connected component of 18 cadmium-AOPs. Further, we analyzed the directed network of 59 KEs and 82 key event relationships (KERs) in the largest component using graph-theoretic approaches. Subsequently, we mined published literature using artificial intelligence-based tools to provide auxiliary evidence of cadmium association for all KEs in the largest component. Finally, we performed case studies to verify the rationality of cadmium-induced toxicity in humans and aquatic species. Overall, cadmium-AOP network constructed in this study will aid ongoing research in systems toxicology and chemical exposome.

## 1. Introduction

Heavy metals are naturally occurring dense elements that are usually toxic in nature [1,2]. The rising levels of heavy metals in the environment, owing to various industrial and anthropogenic activities, is cause for grave concern as they can negatively affect human health and environment [1,3]. Cadmium is one such heavy metal which contaminates both terrestrial and aquatic environments, and is a major contributor of toxicity in various exposomes [4–6]. The prolonged biological half-life of cadmium coupled with its low excretion rates promotes accumulation of cadmium in humans and causes a wide range of disorders [5,7–11]. Similarly, cadmium accumulation in marine organisms, especially fish, has been reported to disrupt the endocrine systems and cause various reproductive and developmental disorders [12–16]. Due to its wide range of toxicities, cadmium has been identified as a priority pollutant by the United States Environmental Protection Agency (US EPA) [17]. Moreover, cadmium is classified as a carcinogen by the International Agency for Research on Cancer (IARC) [18].

A landmark report titled ‘Toxicity testing in the 21st century: a vision and a strategy’ published by the US national research council, recommended the development of conceptual frameworks to enable rapid, efficient and cost-effective characterization of toxic chemicals [19]. Inspired by this report, Ankley *et al.* [20] proposed a conceptual framework that involves aggregation and organization of existing mechanistic data on adverse outcomes induced by chemical exposure, and termed it as adverse outcome pathways (AOPs). Subsequently, the Organisation of Economic Co-operation and Development (OECD) launched an international effort to develop AOPs that can guide chemical risk assessment and regulatory decisions, and setup a user-friendly open-source repository named AOP-Wiki (https://aopwiki.org/) to enable collaborative development and evaluation of AOPs.

AOP is a conceptual toxicological knowledge framework that consists of sequentially ordered biological events underlying a stressor (e.g., chemicals) – induced adverse biological outcome [20,21]. Here, the biological events are organized into modular constructs termed as key events (KEs) and are connected to each other through directional links termed as key event relationships (KERs) [21–23]. In an AOP, the originating KE that involves the molecular interaction between the stressor and the biological target is termed as molecular initiating event (MIE) [24]. The anchored biological events that are at organ or higher level of biological organization and are of regulatory relevance, are termed as adverse outcomes (AOs) [24]. The linear setup of individual AOPs limits its ability to capture the biological complexity and diversity of toxicity pathways induced by a stressor or exhaustively capture all perturbations leading to an adverse biological outcome [25]. To address this knowledge gap, the concept of AOP networks was proposed [25,26]. Construction of AOP networks highlighted the interactions among individual AOPs enabling the understanding of complex toxicity pathways [25–29]. Therefore, to assess inorganic cadmium-induced toxicity, it is imperative to construct an AOP network specific to cadmium and its inorganic compounds.

Till date, AOP-Wiki has been leveraged to build over 32 different AOP networks for a variety of adverse outcomes such as reproductive disorders [30–33], neurologic disorders [34–37], endocrine disorders [25,38–43], developmental disorders [30,40,44,45], hepatic disorders [25,46,47], pulmonary disorders [48–50], and others [26,27,51–58]. Importantly, Chai *et al.* [33] had integrated the data within Comparative Toxicogenomics Database (CTD) [59] and AOP-Wiki to construct an AOP network specific to arsenic-induced reproductive toxicity. Jeong *et al.* [50] had additionally leveraged the chemical, gene, phenotype, and disease associations within CTD to construct an AOP network specific to pulmonary fibrosis. Ravichandran *et al.* [43] had leveraged endocrine-mediated endpoints from DEDuCT [60,61] to construct an AOP network specific to endocrine disruption. Knapen *et al.* [25] had leveraged ToxCast [62] assays to construct an AOP network specific to chemical mixtures in wastewater. In each of the above-mentioned studies, the integration of data from an external source had aided in identification of novel associations with existing AOPs, which resulted in expanded coverage of possible toxicity pathways. Therefore, it is imperative to integrate heterogeneous datasets from various exposome-relevant resources to construct an AOP network relevant for cadmium-induced toxicity.

In this contribution, we aimed to construct and analyze an AOP network specific to inorganic cadmium-induced toxicity. To achieve this, we first extracted the AOP data from AOP-Wiki, systematically assessed their quality and completeness, and retrieved high confidence AOPs. Thereafter, we integrated datasets from five exposome-relevant resources namely, AOP-Wiki, CTD, ToxCast, DEDuCT and NeurotoxKb [63], and identified KEs associated with inorganic cadmium. Subsequently, we leveraged the associated KEs and identified high confidence AOPs that are relevant for cadmium-induced toxicity (cadmium-AOPs). Thereafter, we constructed an AOP network comprising cadmium-AOPs and identified connected components. Further, using graph-theoretic approaches, we analyzed the directed network of the largest component in the cadmium-AOP network. Finally, we leveraged artificial intelligence (AI) based tools to provide auxiliary evidence of cadmium toxicity for each of the KEs present in the largest component. In sum, this is the first study integrating data-centric approaches to construct and characterize an AOP network relevant to cadmium-induced toxicity.

## 2. Methods

### 2.1. Compilation of AOPs from AOP-Wiki

AOP-Wiki (https://aopwiki.org/) is a large public repository hosted by the Society for the Advancement of Adverse Outcome Pathways (SAAOP), and the online resource compiles detailed qualitative information on AOPs that are being developed globally. In AOP-Wiki, AOPs are stored as a set of sequential and measurable key events (KEs) which are linked together by key event relationships (KERs). The KEs capturing molecular responses caused by stressors are termed as molecular initiating events (MIEs). A stressor typically corresponds to chemicals whose exposure can lead to adverse effects. MIEs initiate a cascade of downstream biological responses across different levels of organization such as cellular, tissue, organ and individual. The anchor KEs at organ level of organization or higher, that are generally accepted as being of regulatory significance, are termed as adverse outcomes (AOs) [24,64].

To access the information compiled in AOP-Wiki, we downloaded the latest XML file (released on 1 April 2023) from the AOP-Wiki ‘Projects Downloads’ page (https://aopwiki.org/downloads) which was last accessed on 31 October 2023. Thereafter, we used an in-house python script to parse and retrieve information from the downloaded XML file. For each AOP in the downloaded XML file, we retrieved information on AOP identifier, AOP title, associated KEs (including MIEs and AOs) and KERs, linked stressors, and the status according to OECD and SAAOP. Additionally, we retrieved the biological applicability information for AOPs such as taxonomy, sex and life-stage of the organism, and their weight of evidence. For each KE associated with an AOP, we retrieved the corresponding KE identifier, KE title, level of biological organization, action name, object name and identifiers, and process name, source and identifiers. For each KER associated with an AOP, we retrieved corresponding information on upstream and downstream KEs, adjacency, evidence for biological plausibility, and extent of quantitative understanding. Note that, an adjacency value of ‘Adjacent’ suggests the existence of direct link between the upstream and downstream KEs, whereas a value of ‘Non-adjacent’ suggests the existence of intermediate KE(s) (https://aopwiki.org/handbooks/4).

### 2.2. Filtration of high confidence AOPs within AOP-Wiki

AOP-Wiki is a living document as several AOPs are under development resulting in continuous update and improvement of the resource (https://aopwiki.org/handbooks/4). Therefore, it is crucial to assess the quality and completeness of AOPs including the associated information before considering them to build specific AOP networks [26,43]. Building upon the earlier work by Ravichandran *et al.* [43], we developed a detailed workflow (Figure 1) to filter high confidence AOPs after assessing quality and completeness of AOPs in AOP-Wiki, by employing both computational methods and manual curation efforts in tandem.

**Figure 1:**
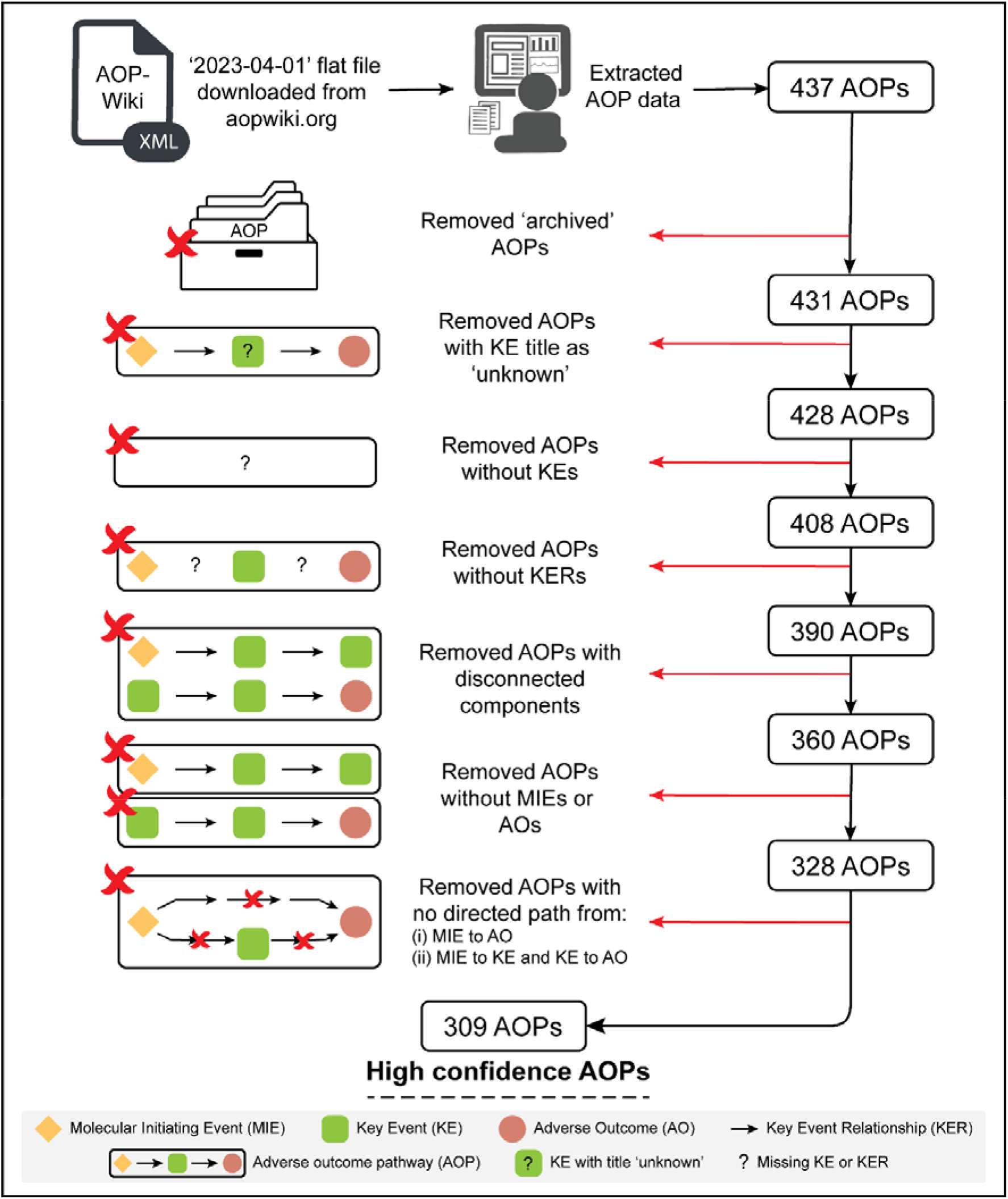
Workflow to filter high confidence adverse outcome pathways (AOPs) from AOP-Wiki by employing computation and manual curation in conjunction.

Initially, using an in-house python script, we retrieved information on 437 AOPs from the downloaded AOP-Wiki XML file. First, we checked the SAAOP status and removed 6 ‘archived’ AOPs (Figure 1), as they are not under active development and are marked as unsuitable for further adoption [65]. Then, we checked and removed 3 AOPs that contained at least one KE with title as ‘unknown’, due to the uncertainty in the associated biological event (Figure 1).

Subsequently, we checked the remaining 428 AOPs to remove empty AOPs which lack KEs. To ensure that the downloaded XML file from AOP-Wiki was up-to-date with information on the AOP page in the online repository, we manually updated any empty AOP determined based on information in the downloaded XML file, if KEs were listed in the ‘Events’ table in the ‘Summary of the AOP’ section of the corresponding AOP page in AOP-Wiki (last accessed on 19 November 2023). Furthermore, for an empty AOP determined based on information in the downloaded XML file, we also checked the ‘Graphical Representation’ section when the ‘Events’ table was empty on the corresponding AOP page in AOP-Wiki (last accessed on 19 November 2023). This combined computational and manual effort led to the removal of 20 empty AOPs which lack KEs (Figure 1).

Next, we checked the remaining 408 AOPs for complete absence of KERs. We manually updated the AOPs lacking KERs based on information in the downloaded XML file, if KERs were listed in the ‘Relationships Between Two Key Events’ table in the ‘Summary of the AOP’ section of the corresponding AOP page in AOP-Wiki (last accessed on 19 November 2023). Furthermore, for an AOP lacking KERs based on information in the downloaded XML file, we also checked the ‘Graphical Representation’ section when the ‘Relationships Between Two Key Events’ table was empty. This combined computational and manual effort led to the removal of 18 AOPs with complete absence of KERs (Figure 1).

Next, we checked for the presence of disconnected components in the remaining 390 AOPs. This computation of the number of connected components in an AOP was performed using the NetworkX [66] python package (https://networkx.org/). We manually updated the AOPs containing disconnected components (i.e., containing more than one connected component) based on information in the downloaded XML file, if additional KERs were available in the ‘Relationships Between Two Key Events’ table in the ‘Summary of the AOP’ section of the corresponding AOP page in AOP-Wiki (last accessed on 19 November 2023). This combined computational and manual effort led to removal of 30 AOPs containing disconnected components (Figure 1).

Subsequently, we checked the remaining 360 AOPs for the presence of at least one MIE and at least one AO (Figure 1). We manually updated the AOPs containing no MIE and/or no AO based on information in the downloaded XML file, if additional information was available on the ‘Events’ table in the ‘Summary of the AOP’ section of the corresponding AOP page in AOP-Wiki (last accessed on 19 November 2023). This led to removal of 32 AOPs lacking MIE and/or AO (Figure 1).

Finally, we checked the remaining 328 AOPs for the existence of: (i) a directed path that originates from at least one MIE and terminates in at least one AO; (ii) a directed path to every KE that originates from at least one MIE; (iii) a directed path from every KE that terminates in at least one AO. We manually updated AOPs that were not complying with all the three path criteria based on information in the downloaded XML file, if additional information was available from the ‘Relationships Between Two Key Events’ table in the ‘Summary of the AOP’ section of the corresponding AOP page in AOP-Wiki (last accessed on 19 November 2023). This led to removal of 19 AOPs based on the three path criteria, and the remaining 309 AOPs were designated as high confidence AOPs (Figure 1).

Supplementary Table S1. contains the list of 309 high confidence AOPs filtered using the above-mentioned criteria in this study. These 309 high confidence AOPs comprise of 1054 unique KEs (Supplementary Table S2) and 1599 unique KERs (Supplementary Table S3). Note that, while filtering for the high confidence AOPs (Figure 1), we had to assign identifiers to certain KEs and KERs that were manually compiled from AOP-Wiki, as the corresponding identifiers were not available in AOP-Wiki.

### 2.3. Identification of KEs associated with inorganic cadmium

The aim of this study is to build and investigate the network of AOPs within AOP-Wiki that are relevant for inorganic cadmium-induced toxicity. To this end, we first identified KEs associated with inorganic cadmium using five different sources namely, AOP-Wiki (https://aopwiki.org/), Comparative Toxicogenomics Database (CTD) [59] (https://ctdbase.org/), ToxCast [62], DEDuCT [60,61] (https://cb.imsc.res.in/deduct/) and NeurotoxKb [63] (https://cb.imsc.res.in/neurotoxkb/) (Figure 2).

**Figure 2:**
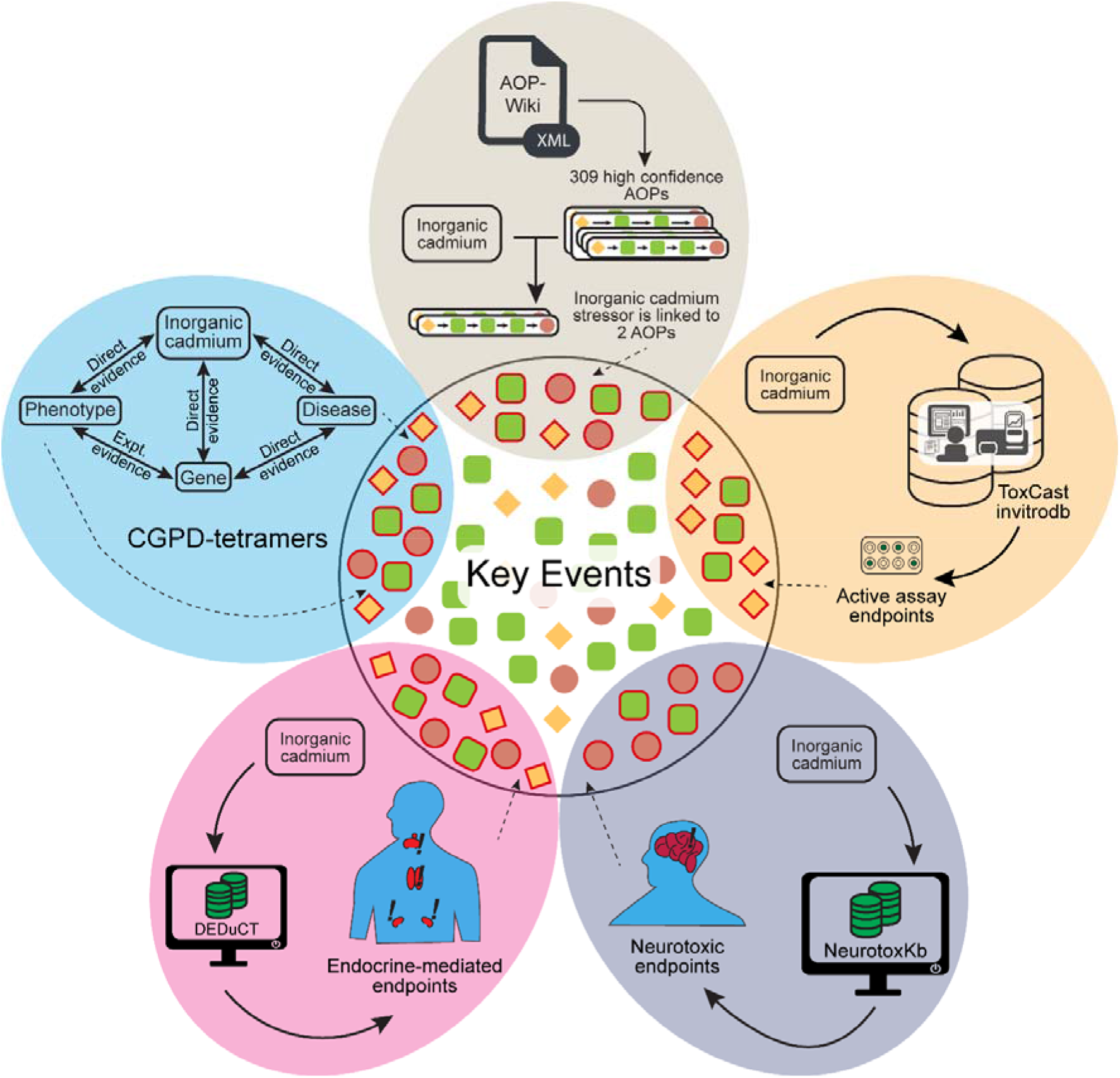
Workflow to identify the key events (KEs) in AOP-Wiki that are associated with inorganic cadmium based on information in five resources namely, AOP-Wiki, CTD, ToxCast, DEDuCT and NeurotoxKb.

#### 2.3.1. KEs associated with inorganic cadmium from AOP-Wiki

For each AOP, we have retrieved information on the stressors that can trigger its progression from the downloaded XML file. We find that 2 of the 309 high confidence AOPs namely, AOP:257 and AOP:296, are documented to be associated with inorganic cadmium, specifically, cadmium and cadmium chloride, in AOP-Wiki. Thus, we considered the 10 KEs comprising the 2 AOPs to be associated with inorganic cadmium (Figure 2).

#### 2.3.2. Identification of KEs associated with inorganic cadmium using CGPD-tetramers from CTD

CTD [59] (https://ctdbase.org/) is among the largest toxicogenomics resources that compiles published information on health effects due to chemical exposures. CTD provides information on associations among chemicals, genes or proteins, pathways, phenotypes, and diseases through systematic curation from published literature and other resources. Recently, Jeong *et al.* [50] leveraged the chemical (C), gene (G), phenotype (P) and disease (D) tetramers, i.e., CGPD-tetramers, constructed from the data compiled in CTD, to identify the KEs associated with pulmonary fibrosis. Here, we followed the workflow proposed by Davis *et al.* [67] to construct the CGPD-tetramers specific to inorganic cadmium, and thereafter, leveraged them to identify the associated KEs in AOP-Wiki.

Specifically, we leveraged data from CTD’s September 2023 release (last accessed on 31 October 2023) to construct the CGPD-tetramers specific to inorganic cadmium. First, we identified the list of chemicals within CTD which correspond to inorganic cadmium, namely, cadmium, cadmium chloride, cadmium nitrate, cadmium oxide, cadmium sulfate, cadmium sulfide, cadmium telluride and cadmium selenide. For each of these chemicals, we compiled the corresponding chemical-gene, chemical-phenotype and chemical-disease, and gene-disease pairs from CTD. Further, we leveraged GO (Gene Ontology) annotations of genes from NCBI Gene resource (https://ftp.ncbi.nih.gov/gene) (last accessed on 31 October 2023) to identify gene-phenotype pairs. Thereafter, to construct the inorganic cadmium specific CGPD-tetramers, we considered: (i) chemical-gene and chemical-phenotype associations with literature evidence; (ii) chemical-disease and gene-disease associations with ‘marker/mechanism’ or ‘marker/mechanism|therapeutic’ evidence; (iii) gene-phenotype associations with GO annotations based on only the experimental results (https://geneontology.org/docs/guide-go-evidence-codes/). This construction procedure resulted in a non-redundant list of 9873 CGPD-tetramers which comprise 3 chemicals (cadmium, cadmium chloride, and cadmium sulfate), 849 genes, 309 phenotypes (GO terms), and 163 disease terms (Supplementary Table S4). Subsequently, we manually mapped the phenotype and disease terms in 9873 CGPD-tetramers specific to inorganic cadmium to the KEs in AOP-Wiki, in order to identify the KEs associated with inorganic cadmium.

For the CGPD-tetramer phenotype linked GO terms, we generated the immediate neighbor terms (both parent and children GO terms) using the GOSim [68] package available in R programming language. Next, we overlapped the GO terms (along with their neighbor terms) with the process identifiers of KEs in AOP-Wiki, and manually inspected the KE title and phenotype name before accepting any mapping between CTD phenotypes and KEs in AOP-Wiki. Through this exercise, we were able to map 88 phenotypes in CGPD-tetramers to 181 KEs in AOP-Wiki (Figure 2). For the CGPD-tetramer disease terms, we used disease identifier and disease name, and manually mapped them to KEs in AOP-Wiki using the process identifiers and the title of KEs. Through this exercise, we were able to map 70 disease terms in CGPD-tetramers to 60 KEs in AOP-Wiki (Figure 2).

#### 2.3.3. Identification of KEs associated with inorganic cadmium using ToxCast assay endpoints

ToxCast is a US EPA project, which has experimentally screened nearly 10000 environmental chemicals to assess their adverse effects. Here, we also leveraged the ToxCast assay endpoints reported for inorganic cadmium to identify the associated KEs in AOP-Wiki. To this end, we downloaded the latest ToxCast invitrodb version 4.1 dataset [69] from the US EPA repository (https://www.epa.gov/chemical-research/exploring-toxcast-data), and thereafter, retrieved the active assay endpoints (‘hitc’ ≥ 0.9) for two inorganic cadmium compounds namely, cadmium chloride and cadmium nitrate, from the ‘mc5-6_winning_model_fits-flags_invitrodb_v4_1_SEPT2023.csv’ file. In ToxCast, the ‘activatory’ or ‘inhibitory’ effect of a chemical is obtained from the sign of the ‘top’ value of the winning model mentioned in the ‘mc4_all_model_fits_invitrodb_v4_1_SEPT2023.csv’ file [70]. Further, we accessed the ‘assay_gene_mappings_invitrodb_v4_1_SEPT2023.xlsx’ file to obtain the biological metadata for the shortlisted assay endpoints.

Subsequently, we manually mapped the shortlisted assay endpoints in ToxCast for the 2 inorganic cadmium compounds to KEs in AOP-Wiki. To elaborate, we first overlapped the target biological process, gene identifier, gene names and gene aliases corresponding to the assay endpoints in ToxCast with the KEs in AOP-Wiki based on their titles, object identifier and object name. Next, we manually inspected the assay endpoint description, ‘activatory’ or ‘inhibitory’ effect of the chemical in the assay endpoint, and the action of the overlapped KE, prior to accepting a mapping between an assay endpoint in ToxCast and KE in AOP-Wiki. This procedure resulted in the mapping of 30 KEs in AOP-Wiki to 28 ToxCast assay endpoints specific to 2 inorganic cadmium compounds (Figure 2).

#### 2.3.4. Identification of KEs associated with inorganic cadmium using DEDuCT and NeurotoxKb

DEDuCT [60,61] is among the largest resources on endocrine disrupting chemicals that has compiled manually curated information on such chemicals along with supporting evidence from published literature. Here, we also leveraged DEDuCT to identify KEs in AOP-Wiki associated with inorganic cadmium. Specifically, we compiled the endocrine-mediated endpoints for three inorganic cadmium compounds namely, cadmium, cadmium chloride, and cadmium nitrate, from DEDuCT (https://cb.imsc.res.in/deduct/), and thereafter, manually mapped the endpoints to KEs in AOP-Wiki based on their titles. This procedure resulted in the mapping of 58 KEs in AOP-Wiki to 33 DEDuCT endpoints specific to 3 inorganic cadmium compounds (Figure 2).

NeurotoxKb [63] is a dedicated resource that has compiled manually curated information on neurotoxicants along with supporting evidence in mammals from published literature. Here, we also leveraged NeurotoxKb to identify KEs in AOP-Wiki associated with inorganic cadmium. Specifically, we compiled the neurotoxic endpoints for two inorganic cadmium compounds namely, cadmium and cadmium chloride, from NeurotoxKb (https://cb.imsc.res.in/neurotoxkb/), and thereafter, manually mapped the endpoints to KEs in AOP-Wiki based on their titles. This procedure resulted in the mapping of 7 KEs in AOP-Wiki to 5 NeurotoxKb endpoints specific to 2 inorganic cadmium compounds (Figure 2).

Overall, by integrating information contained in AOP-Wiki, CTD, ToxCast, DEDuCT and NeurotoxKb, we compiled a list of 312 KEs (Supplementary Table S5) in AOP-Wiki with published evidence of being associated with inorganic cadmium, specifically, cadmium, cadmium chloride, cadmium sulfate and cadmium nitrate.

### 2.4. Curated subset of AOPs relevant for cadmium-induced toxicity

After compiling the list of 312 KEs associated with inorganic cadmium, we find that 241 of the 309 high confidence AOPs contain at least one KE associated with inorganic cadmium (Figure 3). Of these, we find that 34 high confidence AOPs have at least one MIE and at least one AO associated with inorganic cadmium (Figure 3). Moreover, we ascertained that the 34 high confidence AOPs have at least one directed path that originates from a MIE associated with inorganic cadmium and terminates in an AO associated with inorganic cadmium (Figure 3). Lastly, for each of these 34 high confidence AOPs, we computed the coverage score [33] which is the ratio of the number of KEs associated with inorganic cadmium to the total number of KEs in an AOP. By imposing a coverage score threshold of ≥ 0.4 [33], we identified a subset of 30 high confidence AOPs as relevant for cadmium-induced toxicity and designated them as ‘cadmium-AOPs’ (Figure 3). Subsequently, we considered the subset of 30 cadmium-AOPs to construct an AOP network relevant for cadmium-induced toxicity.

**Figure 3:**
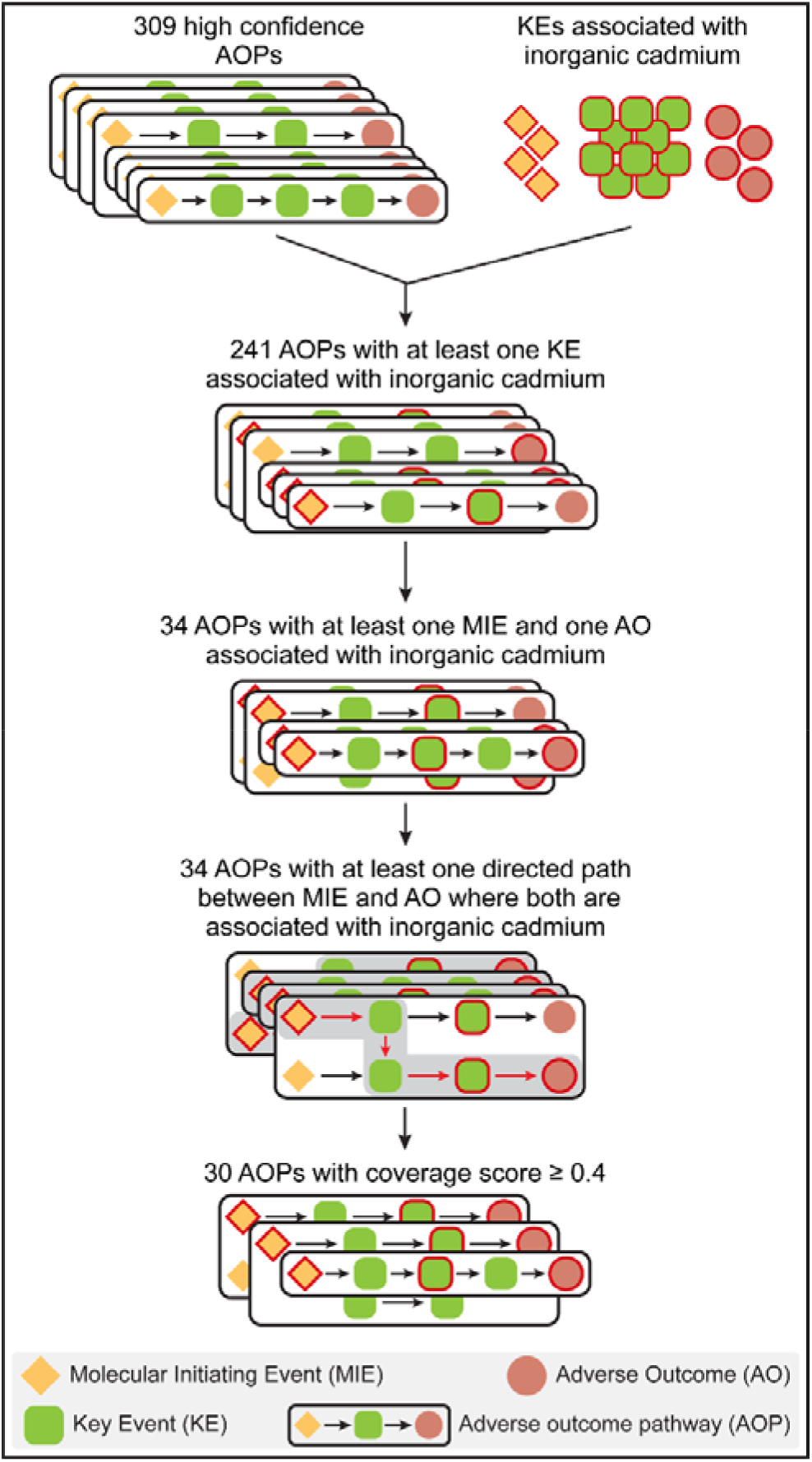
Workflow to identify AOPs relevant for cadmium-induced toxicity, that is, cadmium-AOPs, from the curated list of high confidence AOPs.

The 30 cadmium-AOPs comprise 98 unique KEs and 130 unique KERs. For each KER in an AOP, AOP-Wiki provides corresponding information on the weight of the evidence (WoE) for its biological plausibility (Supplementary Table S6). Following Ravichandran *et al.* [43], we leveraged this WoE information for KERs in terms of ‘High’, ‘Moderate’, ‘Low’ or ‘Not Specified’, to compute the fraction of KERs in an AOP with ‘High’ WoE [i.e., F(High)], the fraction of KERs in an AOP with ‘Moderate’ WoE [i.e., F(Moderate)], the fraction of KERs in an AOP with ‘Low’ WoE [i.e., F(Low)], and the fraction of KERs in an AOP with ‘Not Specified’ WoE [i.e., F(Not Specified)]. Thereafter, we assigned the cumulative WoE using the following criteria [43]:

i. if F(High) ≥ 0.5, the cumulative WoE of the AOP is assigned as ‘High’,
ii. if F(High) < 0.5, but (F(High) + F(Moderate)) ≥ 0.5, the cumulative WoE of the AOP is assigned as ‘Moderate’,
iii. if (F(High) + F(Moderate)) < 0.5, but (F(High) + F(Moderate) + F(Low)) ≥ 0.5, the cumulative WoE of the AOP is assigned as ‘Low’,
iv. if none of the above-mentioned three conditions are satisfied, the cumulative WoE of the AOP is assigned as ‘Not Specified’.

Supplementary Table S7. provides the cumulative WoE for each of the 30 cadmium-AOPs. For each of the 30 cadmium-AOPs, we have also compiled information on taxonomic, sex, and life-stage applicability along with the corresponding levels of evidence (Supplementary Table S7).

### 2.5. AOP network construction and visualization

To better understand the shared relationships among the 30 cadmium-AOPs, we constructed an undirected AOP network based on shared KEs among the AOPs. In the undirected AOP network, nodes correspond to 30 cadmium-AOPs, and there exists an edge between any pair of cadmium-AOPs if they share at least one KE (Figure 4). After constructing the undirected AOP network comprising 30 cadmium-AOPs, we visualized and identified the connected components in the network using Cytoscape [71].

**Figure 4:**
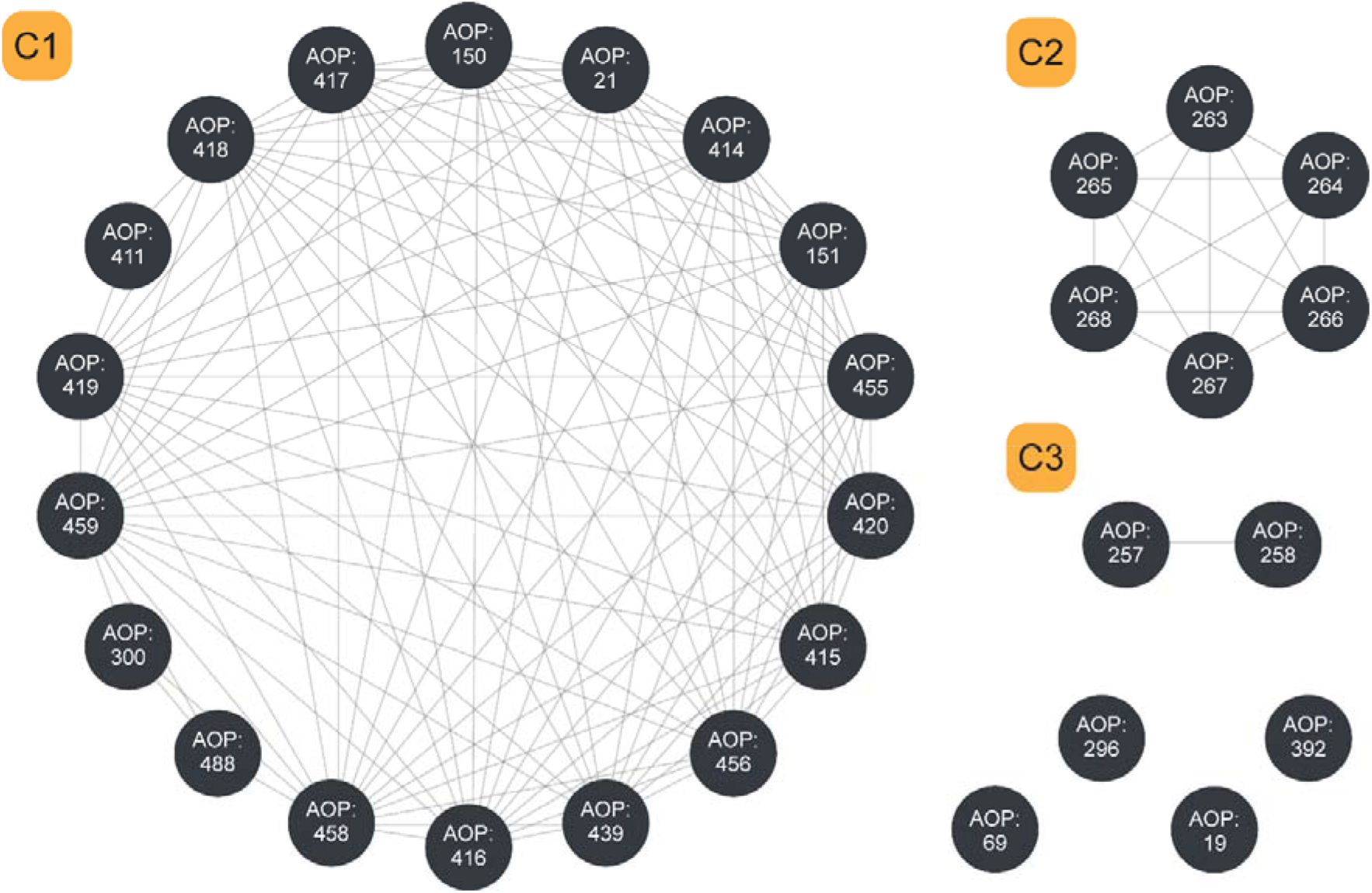
Undirected network of 30 cadmium-AOPs, where each node corresponds to a cadmium-AOP and there exists an edge between any two cadmium-AOPs that share at least one KE. This network has 3 connected components (with two or more nodes) which are labelled as C1, C2, and C3, and 4 isolated nodes.

Moreover, we constructed a directed AOP network comprising KEs and KERs in the 30 cadmium-AOPs. Since the undirected AOP network consists of multiple disconnected components (Figure 4), the directed AOP network also comprises multiple components. In particular, we constructed the directed network corresponding to the largest connected component of the AOP network (Figure 5). In the directed network, the nodes represent KEs and a directed edge represents a KER linking its upstream KE to its downstream KE. For the directed AOP network, we computed different network measures namely, in-degree, out-degree, eccentricity, and betweenness centrality using NetworkX [66] python package.

**Figure 5:**
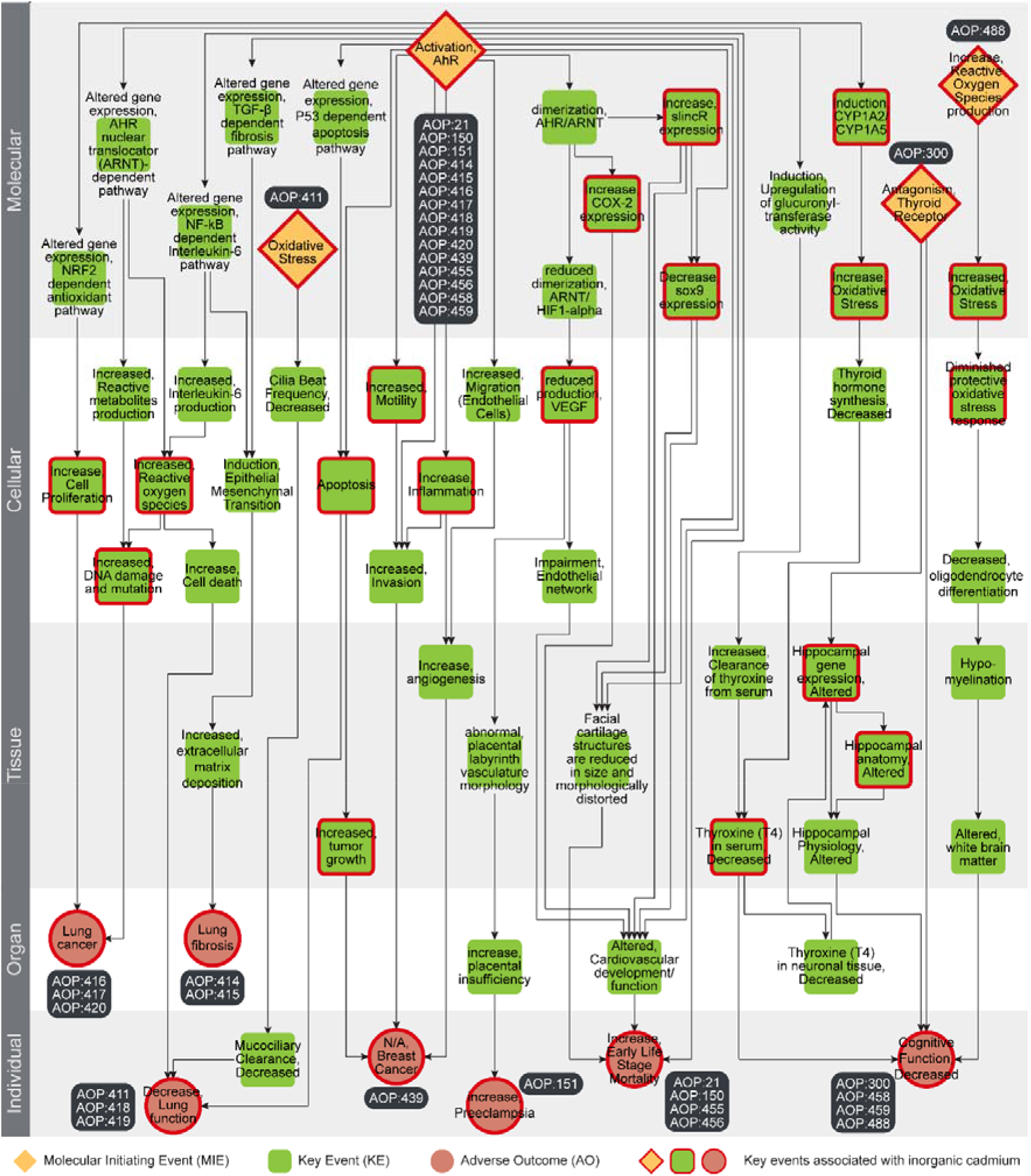
Directed network corresponding to the largest connected component (C1) in the cadmium-AOP network comprising of 59 KEs and 82 KERs. Among the 59 KEs, 4 are categorized as MIEs (denoted as diamond), 7 are categorized as AOs (denoted as circle), and the remaining 48 are categorized as KEs (denoted as rounded square). The 29 KEs (including MIEs and AOs) associated with inorganic cadmium are marked in ‘red’. In this figure, the 59 KEs are arranged vertically according to their level of biological organization.

## 3. Results and Discussion

### 3.1. Integrative data-centric construction and analysis of cadmium-AOP network

In this study, we aimed to construct and analyze an adverse outcome pathway (AOP) network relevant for inorganic cadmium-induced toxicity. To achieve this, we first retrieved the AOP data from AOP-Wiki, assessed their quality and completeness, and curated 309 high confidence AOPs (Methods; Figure 1; Supplementary Table S1). Thereafter, by leveraging various exposome-relevant databases such as AOP-Wiki, CTD, ToxCast, DEDuCT and NeurotoxKb, we systematically identified 312 key events (KEs) present in AOP-Wiki to be associated with inorganic cadmium-induced toxicity (Methods; Figure 2; Supplementary Table S5). Finally, by applying various criteria on the specialized KEs such as molecular initiating events (MIEs) and adverse outcomes (AOs), and considering a coverage score cut-off of 0.4, we identified 30 high confidence AOPs relevant for inorganic cadmium-induced toxicity (which are designated as ‘cadmium-AOPs’) (Methods; Figure 3). Importantly, we emphasize that AOP-Wiki had linked only 2 AOPs (AOP:257 and AOP:296) to inorganic cadmium stressor, whereas our systematic integration of heterogeneous data from diverse sources led to the identification of 28 additional AOPs within AOP-Wiki to be relevant for inorganic cadmium-induced toxicity.

Among the 30 cadmium-AOPs, we observed 26 cadmium-AOPs have a coverage score ≥ 0.5 signifying that at least half of their KEs have published evidence of being associated with inorganic cadmium (Methods; Table 1). Notably, 24 of these 26 cadmium-AOPs with high coverage score are identified through our integrative data-centric approach to be relevant for inorganic cadmium-induced toxicity. Further, we observed 9 cadmium-AOPs have ‘High’ cumulative weight of evidence (WoE) and 6 cadmium-AOPs have ‘Moderate’ cumulative WoE, highlighting the significance of the identified AOPs for inorganic cadmium-induced toxicity (Methods; Table 1). Based on the domain of taxonomic applicability mentioned in AOP-Wiki, we observed 17 out of 30 cadmium-AOPs are applicable across diverse group of species such as humans, animals like rats, mice, and chicken, and aquatic species like zebrafish and *Lemna minor* (Supplementary Table S7). Furthermore, based on the domain of life-stage applicability mentioned in AOP-Wiki, we find that the 30 cadmium-AOPs capture the potential of inorganic cadmium to induce toxicity in various developmental stages (Supplementary Table S7). In addition, this underscores the relevance of the identified cadmium-AOPs in the assessment of inorganic cadmium toxicity in humans and in ecologically relevant species.

**Table 1:**
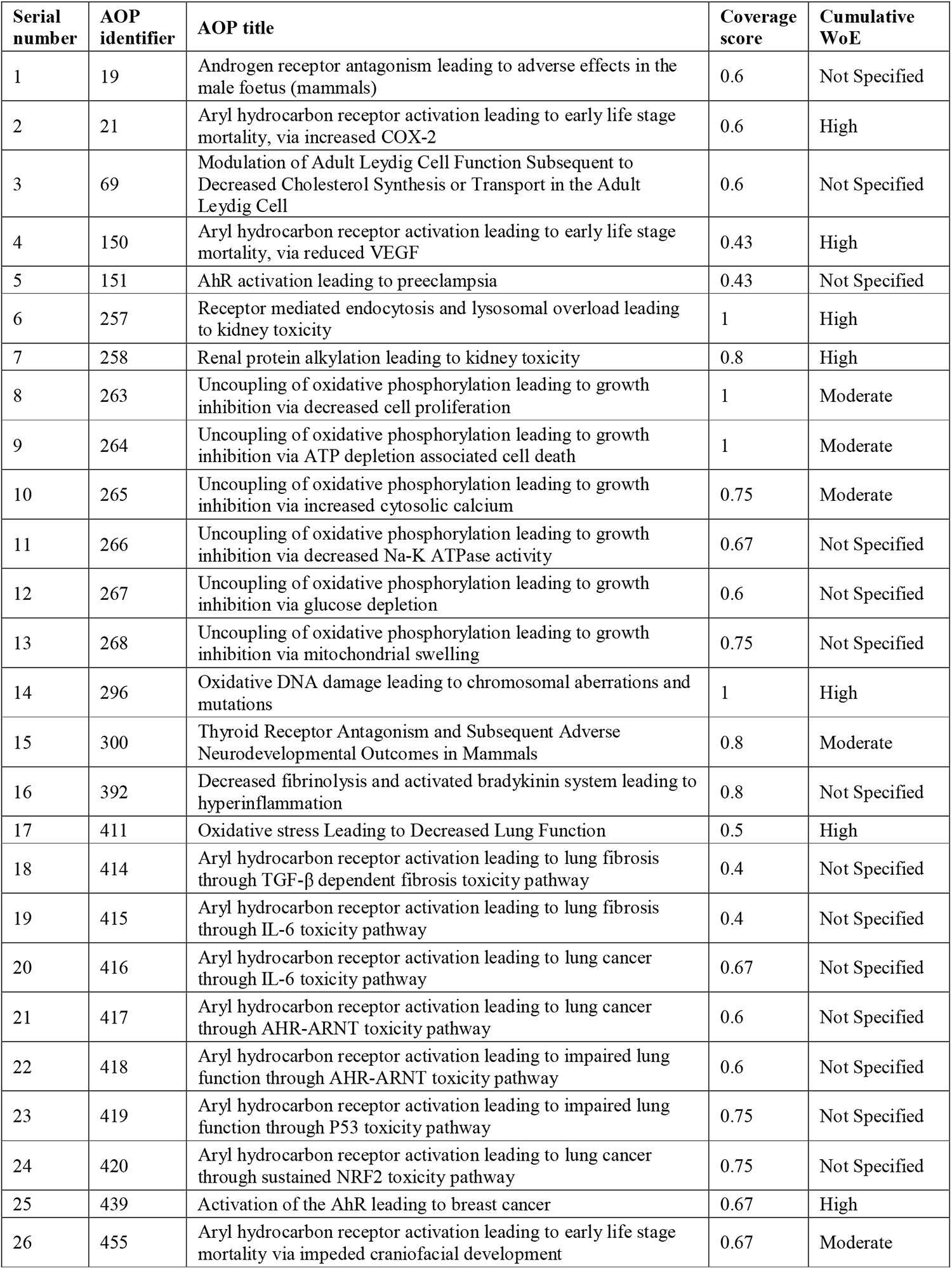
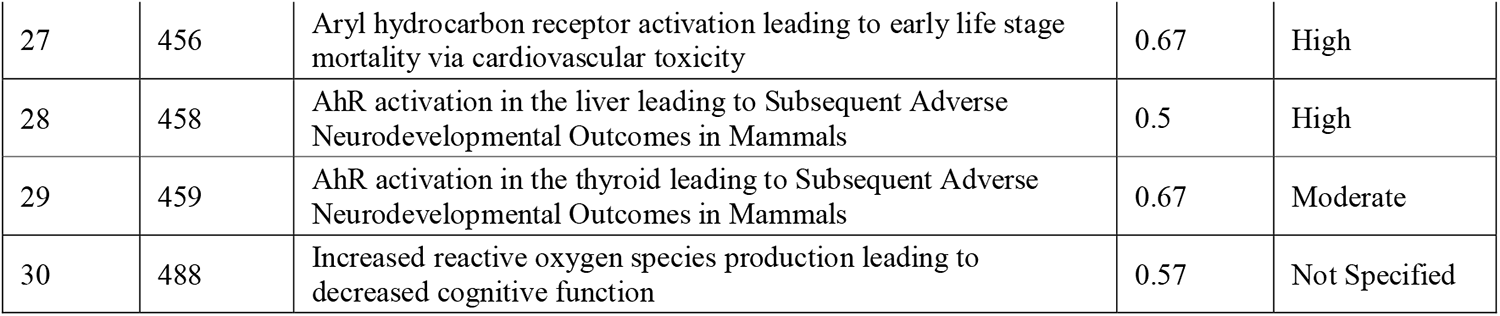
The curated list of 30 cadmium-AOPs and their corresponding AOP identifiers, AOP titles, computed coverage scores, and cumulative weight of evidence (WoE).

Figure 4 shows an undirected network representation of the 30 cadmium-AOPs (i.e., cadmium-AOP network) constructed by considering the cadmium-AOPs as nodes and the existence of shared KEs between any two cadmium-AOPs as edges (Methods). We found 3 connected components with two or more nodes (labeled C1, C2 and C3) and 4 isolated nodes in the cadmium-AOP network. The connected component C1 is the largest connected component (LCC) comprising 18 cadmium-AOPs, followed by C2 comprising 6 cadmium-AOPs, and C3 comprising 2 cadmium-AOPs. We emphasize that only one (AOP:257) of the two cadmium-AOPs linked to inorganic cadmium stressor in AOP-Wiki is part of C3, while the C1 and C2 exclusively comprise cadmium-AOPs identified through our integrative data-centric approach. In other words, all cadmium-AOPs except one that comprise the three clusters C1, C2 and C3 in the cadmium-AOP network were identified to be relevant for inorganic cadmium-induced toxicity by our integrative analysis. Supplementary Table S8 provides the list of cadmium-AOPs, and their corresponding MIEs and AOs associated with each of the connected components.

Among the 18 cadmium-AOPs in LCC, 8 AOPs are related with lung diseases (AOP:411, AOP:414, AOP:415, AOP:416, AOP:417, AOP:418, AOP:419, and AOP:420), 4 AOPs with developmental disorders (AOP:21, AOP:150, AOP:455, and AOP:456), another 4 AOPs with cognitive disorders (AOP:300, AOP:458, AOP:459, and AOP:488), 1 AOP with breast cancer (AOP:439) and 1 AOP with pregnancy disorder namely, preeclampsia (AOP:151) (Supplementary Table S8). ‘Activation, AhR’ (KE:18) is the most common MIE in C1, and is shared across 15 AOPs related to all the adverse outcomes (Supplementary Table S8). The 6 cadmium-AOPs in C2 (AOP:263, AOP:264, AOP:265, AOP:266, AOP:267, and AOP:268) have ‘Decrease, Coupling of oxidative phosphorylation’ (KE:1446) as MIE and ‘Decrease, Growth’ (KE:1521) as AO (Supplementary Table S8). Upon closer inspection, we observed that these AOPs are developed by a single research group to address ecotoxicological effects of stressor induced mitochondrial dysfunction on growth and development of organisms. We remark that these 6 cadmium-AOPs, sharing both MIE and AO, can be considered as variations of a single toxicological pathway and represent potential alternate strategies to understand the same process. The 2 cadmium-AOPs in C3 (AOP:257 and AOP:258) have ‘Occurrence, Kidney toxicity’ (KE:814) as their AO (Supplementary Table S8). Upon closer inspection, we observed that AOP:257 is already documented in AOP-Wiki to be linked to inorganic cadmium stressor, and 4 of the 5 KEs in AOP:258 are associated with inorganic cadmium-induced toxicity.

Furthermore, we observed that C1 comprises 59 unique KEs and 82 unique KERs, C2 comprises 11 unique KEs and 15 unique KERs, and C3 comprises 8 unique KEs and 7 unique KERs. Similar to coverage score of AOPs, we computed the coverage score for each of the connected components based on the fraction of KEs associated with inorganic cadmium (Methods). We observed that C3 had the highest coverage score of 0.88 with 7 of the 8 KEs associated with inorganic cadmium, which can be attributed to the presence of cadmium stressor linked AOP:257. C1 has a coverage score of 0.49 with 29 of 59 KEs associated with inorganic cadmium, and C2 has a coverage score of 0.45 with 5 of 11 KEs associated with inorganic cadmium.

In sum, the undirected cadmium-AOP network highlighted the connectedness of different AOPs relevant for inorganic cadmium-induced toxicity. Connected component C1 is the largest, has roughly half the KEs associated with inorganic cadmium, and is the most diverse in terms of MIEs and AOs. We therefore considered C1 for further network-based analysis in this study.

### 3.2. Characterization and network-based analysis of the largest component in cadmium-AOP network

A directed network representation of connected AOPs, with nodes as KEs and directed edges as KERs, has the potential to elucidate AOP interactions and reveal connections among toxicity pathways [25]. In this study, we visualized the directed network of 18 cadmium-AOPs in LCC (C1), and analyzed the LCC using network measures to elucidate cadmium-induced toxicity pathways. Notably, all the 18 cadmium-AOPs were identified through our integrative data-centric approach. Furthermore, these 18 cadmium-AOPs comprise AOs that are not linked to inorganic cadmium related stressors in AOP-Wiki.

The LCC C1 comprises 59 unique KEs, of which 4 are MIEs and 7 are AOs (Figure 5). We observed that ‘Activation, AhR’ (KE:18) is the most common MIE, and is shared among 15 AOPs (Figure 5; Supplementary Table S8). ‘Increase, Early Life Stage Mortality’ (KE:947) and ‘Cognitive Function, Decreased’ (KE:402) are the most common AOs, and are shared among 4 AOPs each (Figure 5; Supplementary Table S8). Besides MIEs and AOs, ‘dimerization, AHR/ARNT’ (KE:944) is the most common KE, and is shared amongst 5 AOPs (Figure 5; Supplementary Table S8). Furthermore, there are 82 unique KERs in the LCC, of which 8 KERs are labeled as ‘Non-adjacent’ and others are labeled as ‘Adjacent’ (Supplementary Table S6).

We characterized the 59 KEs in LCC based on additional information available in AOP-Wiki. We observed that 39 of the 59 KEs are applicable across a diverse range of taxonomies. We also observed that 35 of 59 KEs are applicable across all stages of development. Moreover, based on the KE titles, we observed incompleteness and duplications among the 59 KEs. For instance, the AO ‘N/A, Breast Cancer’ (KE:1193) has incomplete action information. The 2 KEs, ‘Increase, Oxidative stress’ (KE:1969) and ‘Increased, Oxidative stress’ (KE:1088), have the same underlying process and action but are reported as two different KEs in AOP-Wiki. Further, we noted that while AOP-Wiki provides stressor information for each AOP, it does not provide associations between stressors and KEs. Therefore, by following a systematic workflow (Methods; Figure 2), we identified 29 of the 59 KEs to be associated with inorganic cadmium-induced toxicity by leveraging compiled information in five resources namely, AOP-Wiki, CTD, ToxCast, DEDuCT and NeurotoxKb (Figure 5).

Furthermore, we computed different node-centric network measures (in-degree, out-degree, eccentricity, betweenness centrality, convergence and divergence) for the constructed directed network of 18 cadmium-AOPs (Methods; Supplementary Table S9). We observed that KE ‘Altered, Cardiovascular development/function’ (KE:317) has the maximum in-degree of 5, while the MIE ‘Activation, AhR’ (KE:18) has the maximum out-degree of 17. Additionally, based on the in-degree and out-degree values of each KE, we identified 14 convergent (i.e., in-degree > out-degree) KEs and 10 divergent (i.e., in-degree < out-degree) KEs (Methods; Supplementary Table S9). Among the convergent KEs, we observed that ‘Altered, Cardiovascular development/function’ (KE:317) has the maximum in-degree value of 5. This KE links different toxicity pathways that originate from activation of AhR and lead to early life-stage mortality (Figure 5). The convergent AO, ‘Cognitive Function, Decreased’, with in-degree value of 4, is the anchor of different toxicity pathways originating from MIEs such as ‘Activation, AhR’ (KE:18), ‘Antagonism, Thyroid Receptor’ (KE:1656), and ‘Increase, Reactive Oxygen Species production’ (KE:257) (Figure 5; Supplementary Table S8). Among the divergent KEs, ‘Activation, AhR’ (KE:18) with the maximum out-degree of 17, is the origin of different toxicity pathways leading to the 7 AOs ‘Lung cancer’ (KE:1670), ‘Lung fibrosis’ (KE:1276), ‘Decrease, Lung function’ (KE:1250), ‘N/A, Breast Cancer’ (KE:1193), ‘Increase, Preeclampsia’ (KE:1893), ‘Increase, Early Life Stage Mortality’ (KE:947), and ‘Cognitive Function, Decreased’ (KE:402) (Figure 5; Supplementary Table S8).

Eccentricity of a node indicates the distance of a node to all the other nodes in the network [72]. A large eccentricity value denotes remotely positioned nodes, whereas the low eccentricity value denotes a more centrally positioned node [72]. We computed the eccentricity of each KE present in the directed AOP network, and observed that 2 MIEs namely, ‘Activation, AhR’ (KE:18), ‘Increase, Reactive Oxygen Species production’ (KE:257), and one KE, ‘Induction, CYP1A2/CYP1A5’ (KE:850) have the maximum eccentricity value of 6 (Supplementary Figure S1; Supplementary Table S9).

Betweenness centrality of a node indicates the proportion of the shortest paths that pass through it to the total number of shortest paths present between all pairs of nodes excluding that node in the network [26]. We computed the betweenness centrality for each of the 59 KEs in the directed AOP network, and observed that ‘Thyroxine (T4) in serum, Decreased’ (KE:281) has the highest value (Supplementary Figure S2; Supplementary Table S9). This KE is present in 2 different toxicity pathways and serves as a critical control event [26] in induced activation of AhR leading to decreased cognitive function.

In sum, the directed AOP network (for LCC) highlighted the diversity of the interconnected inorganic cadmium-induced toxicity pathways. Further, a detailed network analysis highlighted the role of KEs across different toxicity pathways. In the following section, we compile auxiliary evidence and provide detailed explanation of novel association of KEs in LCC with inorganic cadmium-induced toxicity.

### 3.3. Auxiliary evidence for cadmium-induced toxicity pathways in the largest component

In this study, we systematically integrated heterogeneous datasets from AOP-Wiki, CTD, ToxCast, DEDuCT, and NeurotoxKb to identify KEs associated with inorganic cadmium. This data-centric approach enabled us to identify 29 of the 59 KEs present in the directed network of LCC (C1) to be associated with inorganic cadmium (Methods; Figure 5). Notably, the 29 KEs associated with inorganic cadmium comprise 4 MIEs and 7 AOs (Figure 5). The toxicity pathway originating from MIE ‘Activation, AhR’ (KE:18), passing through KEs ‘Apoptosis’ (KE:1262) and ‘Increased, tumor growth’ (KE:1971), and eventually terminating at AO ‘N/A, Breast Cancer’ (KE:1193) contains 4 of the 29 KEs associated with inorganic cadmium. Additionally, we noted that this toxicity pathway is part of AOP ‘Activation of the AhR leading to breast cancer’ (AOP:439), which was systematically developed through extensive literature review [73]. Moreover, AOP:439 has a cumulative WoE of ‘High’ (Table 1) and is applicable to human adults with high level of evidence (Supplementary Table S7). Therefore, the pathway originating from activation of AhR and terminating in breast cancer is a potential toxicity pathway of cadmium-induced breast cancer outcome in humans.

Subsequently, to assess the rationality of associations between the remaining 30 KEs in LCC and inorganic cadmium, we relied on the toxicity data in published literature. We leveraged an artificial intelligence (AI) based tool, AOP-helpFinder [74,75] (https://aop-helpfinder.u-paris-sciences.fr/) to screen existing literature and identify associations of KEs with inorganic cadmium. In addition, we relied on Abstract Sifter [76] (https://comptox.epa.gov/dashboard/chemical/pubmed-abstract-sifter/) to filter published literature from PubMed (https://pubmed.ncbi.nlm.nih.gov/) that are relevant for cadmium-induced toxicity. This extensive literature curation helped us identify novel associations between the remaining 30 KEs in LCC and inorganic cadmium (Supplementary Table S10). We also identified auxiliary evidence for the 29 KEs in LCC that were already associated with inorganic cadmium through our data-centric approach (Supplementary Table S10).

To conclude, we performed two case studies pertaining to a human relevant AOP and ecotoxicity relevant AOPs in LCC to explore the rationale and highlight the relevance of the directed AOP network for cadmium-induced toxicity.

#### 3.3.1 AOP linking inorganic cadmium exposure to preeclampsia

Preeclampsia is a chronic human pregnancy complication, and is emerging as a leading cause of neonatal mortality [77]. We noted that a preeclampsia specific AOP in AOP-Wiki, ‘AhR activation leading to preeclampsia’ (AOP:151), is identified by this study as cadmium-AOP (Table 1) and is part of LCC (C1) (Supplementary Table S8). According to AOP-Wiki, AOP:151 is currently under development, but is included in the OECD work plan (Supplementary Table S1). Different published studies have found significant correlation between environmental exposure to cadmium and preeclampsia in pregnant women [78,79]. Therefore, we leveraged AOP:151 to explore and verify the rationale behind the cadmium-induced toxicity in preeclampsia.

It has been shown that cadmium exposure causes modulation in AhR downstream genes through cross-talk between AhR and estrogen receptors in rat uterine tissue [80]. AhR is a cytosolic protein, which upon binding with a ligand relocates into the nucleus, where it dimerizes with aryl hydrocarbon receptor nuclear translocator (ARNT) to transcribe its downstream genes [81]. Simultaneously, cadmium has been observed to degrade the activity of hypoxia inducible factor 1 (HIF-1) protein, thereby hindering the dimerization of ARNT with HIF-1 [82]. HIF-1 is a master regulator of hypoxia induced responses, and upon dimerization, induces downstream genes such as vascular endothelial growth factor (VEGF) that enables angiogenesis [83,84]. It has been observed that cadmium exposure leads to reduction in VEGF levels of human placental trophoblasts [85] and in human umbilical vein endothelial cells (HUVEC) [86]. Several *in vivo* experiments showed that cadmium exposed pregnant rats have reduced levels of vasculature in placenta, resulting in placental insufficiency [85,87–89]. Alternatively, cadmium exposure has also been seen to cause placental insufficiency by inducing oxidative stress in placenta [90]. Such abrupt vascularization ultimately results in showing key features of preeclampsia in both pregnant rats [91] and in human cell line studies [92]. In conclusion, by leveraging published evidence of cadmium-induced toxicity, we were able to explore a potential toxic pathway in AOP:151 that links cadmium exposure to preeclampsia.

#### 3.3.2 AOPs linking inorganic cadmium exposure to aquatic ecotoxicity

Aquatic ecotoxicity is of primary regulatory concern as it is one of the major determinants in the well-being of terrestrial and aquatic species alike [93]. We identified 2 cadmium-AOPs namely ‘Aryl hydrocarbon receptor activation leading to early life stage mortality, via increased COX-2’ (AOP:21) and ‘Aryl hydrocarbon receptor activation leading to early life stage mortality, via reduced VEGF’ (AOP:150), that are part of LCC (C1), and have ‘High’ evidence of applicability in aquatic species (Supplementary Table S7). Moreover, these AOPs have a cumulative WoE of ‘High’ and are endorsed by Working Group of the National Coordinators of the Test Guidelines Programme (WNT) and the Working Party on Hazard Assessment (WPHA) under the OECD AOP development programme (Supplementary Table S1). Additionally, these 2 AOPs share the same MIE (‘Activation, AhR’) and AO (‘Increase, Early Life Stage Mortality’). AOP:150 also shares 4 of its KEs (including MIE) with AOP:151 discussed above (Figure 5). Therefore, we leveraged these 2 AOPs to explore and verify the rationale behind cadmium toxicity in aquatic ecosystems.

Zebrafish larvae showed dose-dependent response to cadmium toxicity through the upregulation of AhR downstream genes [94]. In zebrafish, the AhR gets activated upon being bound to a ligand, and is transported into the nucleus where it dimerizes with ARNT to enable the transcription of downstream genes [95]. One such group of downstream genes, cyclooxygenase-2 (COX-2) has been observed to be upregulated in common carp spleens upon exposure to inorganic cadmium [96]. Alternatively, cadmium exposure affects HIF-1 activity through AhR mediated pathways, and this results in reduced levels of VEGF [82,85,86]. Various *in vitro* experiments showed that cadmium exposure impairs endothelial cell function and promotes their apoptosis [97–99]. Consequently, cadmium exposure has also been observed to induce cardiovascular developmental disorders by hindering the process of cardiomyocyte differentiation [100,101]. Ultimately, cadmium exposure has been observed to promote early life-stage mortality in aquatic species by hindering their developmental processes [94,102].

## 4. Conclusion

Cadmium, a heavy metal, is considered to be a priority environmental pollutant due to its abundance and considerable toxicity to humans and aquatic species. In the past, the concept of AOP network had enabled elucidation of complex toxicity pathways and aided in regulatory decision making. In this study, we therefore constructed and analyzed an AOP network relevant for inorganic cadmium-induced toxicity, which is expected to aid in regulation of cadmium and its inorganic compounds in future. To construct the AOP network, we first extracted AOPs from AOP-Wiki and systematically curated 309 high confidence AOPs. Simultaneously, we leveraged 5 exposome-relevant resources namely, AOP-Wiki, CTD, ToxCast, DEDuCT, and NeurotoxKb, and integrated the heterogeneous data to identify 312 KEs present in AOP-Wiki to be associated with inorganic cadmium. Subsequently, we integrated the cadmium associated KEs with high confidence AOPs and identified 30 AOPs relevant for cadmium-induced toxicity (cadmium-AOPs). Here, we stress that our integrative data-centric approach revealed 28 novel cadmium-AOPs which were not otherwise linked to cadmium stressor in AOP-Wiki. Thereafter, we constructed the AOP network using the 30 cadmium-AOPs and identified 3 connected components, with the largest component containing 18 cadmium-AOPs. We employed graph-theoretic approaches to analyze the 59 unique KEs present in the largest component and observed that the cadmium-induced MIE, ‘Activation, AhR’ (KE:18), leads to every AO present. Finally, we leveraged an AI based tool namely AOP-helpFinder, and Abstract Sifter to curate supporting evidence for cadmium associations with each of the KEs present in the largest component.

However, we focussed only on AOPs from AOP-Wiki to construct the cadmium-AOP network. Among the 18 cadmium-AOPs within the largest component, only 2 AOPs (AOP:21 and AOP:150) have been endorsed by OECD. We also observed that 9 of these 18 cadmium-AOPs do not compile any evidence for their KERs. Upon closer inspection, we noted that some of the KEs lacked action information or were duplicated, and some KERs directly linked MIE to AO. This can be attributed to the fact that many of these AOPs are under development. Furthermore, we observed that only 66 of the 163 disease terms from CTD associated with cadmium toxicity were mapped to KEs within AOP-Wiki. This highlights that the present cadmium-AOP network may not exhaustively capture the adverse outcomes induced by cadmium toxicity.

Nonetheless, we present the first ever and most up-to-date AOP network specific to cadmium toxicity that is constructed using AOP-Wiki. Our integrative data-centric approach helped in identifying KEs (including MIEs and AOs) associated with inorganic cadmium, which were otherwise not documented within AOP-Wiki. We additionally provide auxiliary evidence for the association of KEs with inorganic cadmium in the directed AOP network. In future, integrative data-centric evidence-based approaches can be leveraged to expand the scope of existing AOPs, or develop novel AOPs, which can further enable the derivation of comprehensive and elaborate AOP networks [103–107]. Further experimentation is required to characterize the points-of-departure of cadmium toxicity and that will aid in strengthening chemical regulations, thereby advancing chemical exposome research.

## Data availability

The data associated with this study is contained in the article or in the supplementary material.

## Acknowledgements

Ajaya Kumar Sahoo would like to thank Ajay Subbaroyan for discussions. Nikhil Chivukula would like to thank Madison Feshuk, Jason Brown and Katie Paul Friedman for their help in accessing ToxCast invitrodb version 4.1 datasets. Areejit Samal would like to acknowledge funding from the Department of Atomic Energy (DAE), Government of India via Apex project to The Institute of Mathematical Sciences (IMSc). The funders have no role in study design, data collection, data analysis, manuscript preparation or decision to publish.

## CRediT author contribution statement

**Ajaya Kumar Sahoo:** Conceptualization, Data Compilation, Data Curation, Formal Analysis, Software, Visualization, Writing; **Nikhil Chivukula:** Conceptualization, Data Compilation, Data Curation, Formal Analysis, Software, Visualization, Writing; **Kundhanathan Ramesh:** Data Compilation, Data Curation, Formal Analysis, Visualization, Writing; **Jasmine Singha:** Formal Analysis, Writing; **Shambanagouda Rudragouda Marigoudar:** Formal Analysis, Writing; **Krishna Venkatarama Sharma:** Formal Analysis, Writing; **Areejit Samal:** Conceptualization, Supervision, Formal Analysis, Writing.

