## Supplementary Figure for "An integrative data-centric approach to derivation and characterization of an adverse outcome pathway network for cadmium-induced toxicity"

### Supplementary Figures S1-S2

### for

#### An integrative data-centric approach to derivation and characterization of an adverse outcome pathway network for cadmium-induced toxicity

Ajaya Kumar Sahoo[a,b,1], Nikhil Chivukula[a,b,1], Kundhanathan Ramesh[a], Jasmine Singha[c], Shambanagouda Rudragouda Marigoudar[c], Krishna Venkatarama Sharma[c], Areejit Samal[a,b,*]

[a] *The Institute of Mathematical Sciences (IMSc), Chennai, India*

[b] *Homi Bhabha National Institute (HBNI), Mumbai, India*

[c] *National Centre for Coastal Research, Ministry of Earth Sciences, Government of India, Pallikaranai, Chennai, India*

[1] A.K.S. and N.C. contributed equally to this work and should be considered as Joint-First authors.

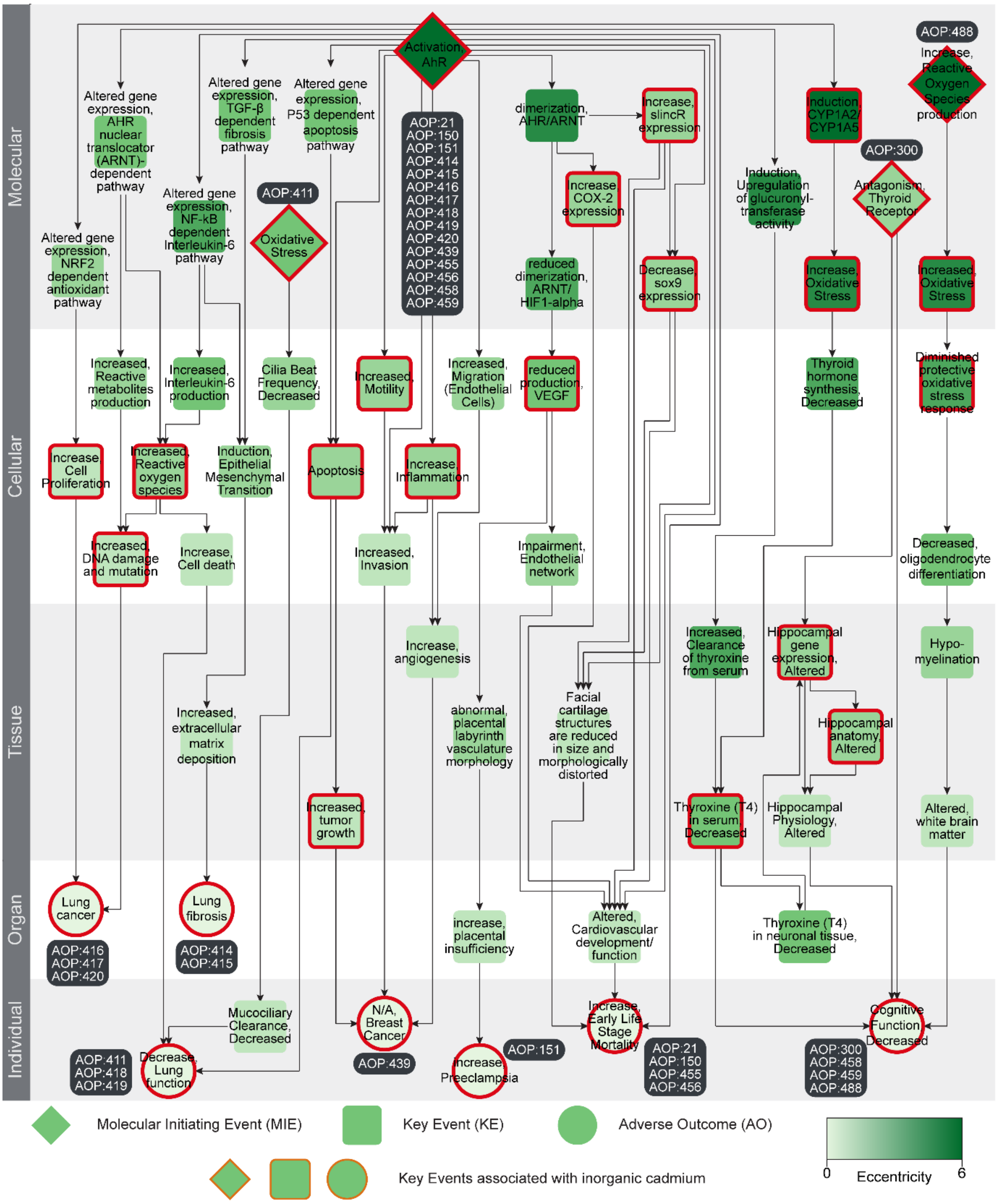


**Figure S1:** Directed network corresponding to the largest connected component (C1) in the cadmium-AOP network, where the KEs (including MIEs and AOs) are colored based on their eccentricity values. The 29 KEs (including MIEs and AOs) associated with inorganic cadmium are marked in 'red'. In this figure, the 59 KEs are arranged vertically according to their level of biological organization.

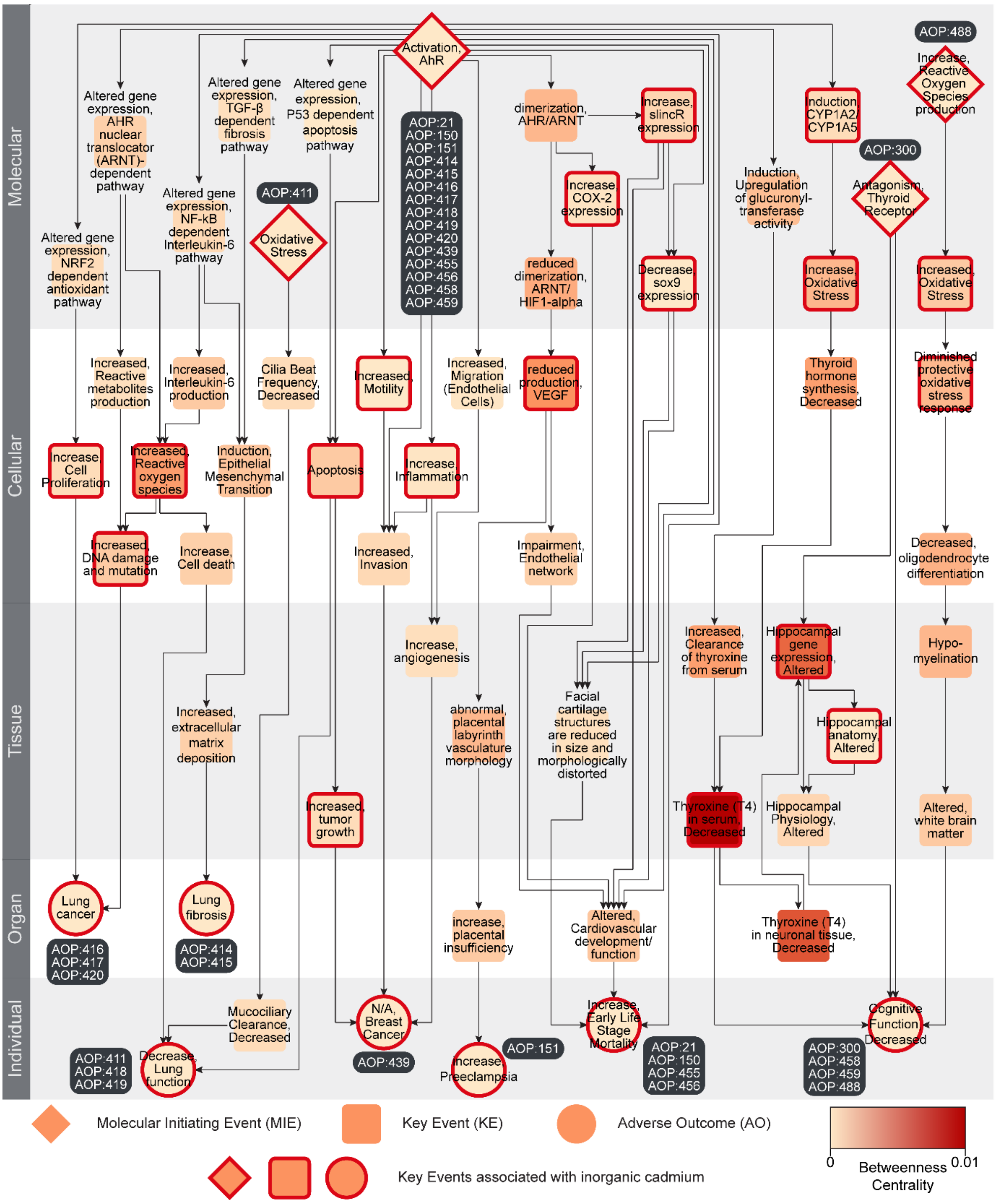


**Figure S2:** Directed network corresponding to the largest connected component (C1) in the cadmium-AOP network, where the KEs (including MIEs and AOs) are colored based on their betweenness centrality values. The 29 KEs (including MIEs and AOs) associated with inorganic cadmium are marked in 'red'. In this figure, the 59 KEs are arranged vertically according to their level of biological organization.
